# Using computational simulations to quantify genetic load and predict extinction risk

**DOI:** 10.1101/2022.08.12.503792

**Authors:** Christopher C. Kyriazis, Jacqueline A. Robinson, Kirk E. Lohmueller

## Abstract

Small and isolated wildlife populations face numerous threats to extinction, among which is the deterioration of fitness due to an accumulation of deleterious genetic variation. Genomic tools are increasingly used to quantify the impacts of deleterious variation in small populations; however, these approaches remain limited by an inability to accurately predict the selective and dominance effects of individual mutations. Computational simulations of deleterious genetic variation offer an alternative and complementary tool that can help overcome these limitations, though such approaches have yet to be widely employed. In this Perspective, we aim to encourage conservation genomics researchers to adopt greater use of computational simulations to aid in quantifying and predicting the threat that deleterious genetic variation poses to extinction. We first provide an overview of the components of a simulation of deleterious genetic variation, describing the key parameters involved in such models. Next, we clarify several misconceptions about an essential simulation parameter, the distribution of fitness effects (DFE) of new mutations, and review recent debates over what the most appropriate DFE parameters are. We conclude by comparing modern simulation tools to those that have long been employed in population viability analysis, weighing the pros and cons of a ‘genomics-informed’ simulation approach, and discussing key areas for future research. Our aim is that this Perspective will facilitate broader use of computational simulations in conservation genomics, enabling a deeper understanding of the threat that deleterious genetic variation poses to biodiversity.

## Introduction

Anthropogenic pressures are resulting in a growing number of small and isolated populations facing an elevated risk of extinction due in part to deleterious genetic factors. Deleterious genetic variation can contribute to extinction in small populations via two related mechanisms: fixation of weakly deleterious alleles due to relaxed negative selection (1, 2) and inbreeding depression due to the exposure of recessive deleterious variation (3, 4). The burden of deleterious variation carried by a population is typically referred to as its “genetic load”, often defined as the reduction in fitness due to segregating and fixed deleterious mutations (4–6).

Genomic tools are now commonly used to quantify deleterious variation and genetic load in wild populations (7–9), though the best approaches for leveraging such datasets to help conserve small populations remains an active area of research. In particular, empirical measures of putatively deleterious variation have seen increased use in conservation genomics studies (9), however, these measures remain relatively crude and often challenging to interpret (10–13).

In light of the limitations of empirical measures of deleterious variation and genetic load, the aim of this review is to encourage more conservation genomics researchers to employ computational genetic simulations and clarify many ongoing debates regarding deleterious mutation parameters. To that end, we first provide an overview of simulations of deleterious genetic variation, illustrating how such approaches can be used to estimate genetic load. Next, we review recent debates on deleterious mutation parameters, focusing on the distribution of fitness effects of new mutations (or DFE), aiming to determine which parameters are best supported by empirical evidence. Finally, we compare modern simulation approaches for modelling inbreeding depression to existing methods that have long been employed in population viability analysis, discussing the pros and cons of “genomics-informed” models of inbreeding depression. Our hope is that this review will provide useful information for researchers aiming to incorporate simulation-based approaches into genomics-based studies of genetic load, enabling more comprehensive assessments of the genomic risks to extinction in small and isolated populations.

### Defining genetic load

Understanding the implications of genetic load for organismal fitness and population viability is a topic of long-standing interest in population and conservation genetics (4–6, 14–17). Several definitions of genetic load have been put forth in the literature recently, often with the aim of partitioning genetic load into “realized” and “potential” load (e.g., (9, 18)). Here, we adhere to the definition of genetic load as the realized reduction in mean fitness in a population due to segregating and fixed deleterious mutations (note that the “mutation load” typically refers only mutations segregating under mutation-selection balance; (5, 6)). The genetic load (L) of a population at a single locus is given by:

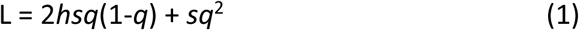

where *s* is the selection coefficient of a mutation, *h* is the dominance coefficient, and *q* is the mutation frequency. Here, the effect of deleterious mutations found as heterozygotes is captured by the 2*hsq*(1-*q*) term and the effect of homozygous deleterious mutations is captured by the *sq*^2^ term. For fixed mutations (*q*=1.0), the genetic load is therefore equal to *s*. Genetic load at a single locus can be related to mean population fitness 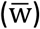 as: 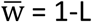. When assuming fitness is multiplicative across sites (i.e., ignoring epistasis and linkage disequilibrium), the mean genome-wide fitness of a population can be obtained by multiplying 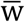 across sites.

This quantity then can be subtracted from 1 to obtain the genome-wide genetic load (6). Thus, the units of genetic load are in terms of multiplicative fitness scaled from 0 to 1.

Another important quantity for predicting the impacts of inbreeding and small population size is the inbreeding load, which quantifies the potential reduction in fitness after inbreeding exposes segregating recessive deleterious mutations (4, 15). Unlike the genetic load, the inbreeding load is measured in terms of lethal equivalents, which represent a summed quantity of *s* for recessive deleterious mutations that are masked as heterozygotes. For a population at equilibrium, the inbreeding load (B) at a single locus is given by (15):

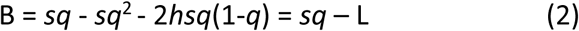

This equation demonstrates that the inbreeding load at a single locus is determined by the frequency and fitness effect of a mutation (*sq*), minus the effects that are expressed as homozygotes (*sq*^2^) and heterozygotes (2*hsq*(1-*q*)). To calculate the total inbreeding load across a diploid genome (2B), this quantity can be summed across sites with deleterious mutations and multiplied by two to account for diploidy.

These fundamental principles demonstrate that an essential component of estimating the genetic load and inbreeding load (hereafter, referred to together as “load”) using genetic variation data is knowing *s* and *h* for individual mutations. However, although some progress has been made in predicting whether a mutation is likely to be neutral or deleterious (e.g., (19–23)), accurately predicting *h* and *s* for individual mutations in genomic sequencing data remains a major challenge, even in humans and model organisms (10–13). For example, a recent simulation study demonstrated that Genomic Evolutionary Rate Profiling (GERP; (22)), a popular method for predicting the deleterious effect of mutations based on evolutionary conservation, cannot reliably distinguish weakly deleterious mutations from strongly deleterious mutations (10), though it is commonly used for this purpose (e.g., (24–27)). Similarly, experimental studies in yeast have found that methods such as SIFT (19) and PROVEAN (23) are poor predictors of the functional impact of a mutation (12, 13) that provide only crude proxies of *s*. Moreover, these methods do not provide any information on dominance, an essential component of quantifying load. These limitations are unlikely to be fully overcome, particularly for non-model organisms, implying that methods for quantifying load based on sequence data will remain somewhat crude approximations for the foreseeable future.

### Overview of simulation-based approaches

Computational simulations using evolutionary models provide an alternate way of quantifying load that alleviates many of the limitations discussed above. Simulations are widely used in population genetics (e.g., (25, 28–34)), yet remain underused in conservation genomics. Historically, this may be due to a relative lack of simulation tools capable of modelling ecologically-realistic scenarios and an often steep learning curve for using simulation software that may be poorly documented (35). However, many of these challenges have been addressed by the forward-in-time genetic simulation program SLiM (36, 37), which offers a flexible array of models incorporating realistic ecological dynamics as well as comprehensive documentation and a graphical user interface. Other similar programs include Nemo (38, 39) and SimBit (40), both of which have been applied in a conservation genetics context (41, 42).

Simulations are broadly useful in evolutionary genetics because they can serve the critical function of revealing which evolutionary scenarios are consistent with observed patterns of genetic variation. All sequence-based evolutionary genetics studies suffer from the limitation that they are observing a single outcome of a stochastic evolutionary process, where underlying mechanisms are largely unobservable. Simulations allow researchers to model this evolutionary process and determine which mechanisms (e.g., genetic drift, gene flow, selection, migration) are needed to explain observed patterns of variation in a dataset. Moreover, the process of using simulations can be extremely valuable for developing intuition on how various evolutionary forces interact to influence patterns of genetic variation, improving the ability of researchers to design evolutionary genetics studies and interpret their results.

For studies aiming to characterize the impacts of small population size on deleterious variation, simulations can be especially useful for quantifying load, which can be directly tabulated from the simulation output (see Supplementary Appendix 1). Simulations can therefore be used to complement empirical measures of load, providing a framework in which to interpret observed patterns and verify that they are expected under a plausible evolutionary model. Moreover, simulations can go beyond empirical measures by projecting load under various future scenarios, illuminating how management actions in the present-day may impact load decades or centuries from now. Finally, modern simulation tools, such as the ecologically-realistic models supported by SLiM3 (36), also offer the potential to conduct an analysis of future extinction risk while incorporating genome-scale genetic variation, analogous to the population viability analysis (PVA) approaches that have long been employed in conservation genetics (e.g., (43–46); see below for further discussion).

In summary, simulation-based approaches have much to offer for genomic studies of deleterious variation in wild populations, yet their application remains relatively limited. In Table 1, we have summarized existing studies that employ simulations along with genomic analyses to investigate genetic load in organisms ranging from ibex to Chinese crocodile lizards.

**Table 1:** Recent studies combining simulations with empirical genomic data to explore impact of small population size on deleterious variation in non-human species.

| Paper | Organism | Simulation software | Distribution of fitness effects | Question addressed with simulations |
| --- | --- | --- | --- | --- |
| Beichman et al. 2022 | Sea otter | SLiM | Kim et al. 2017<sup>^</sup> | How has the fur trade bottleneck impacted genetic load in the sea otter? |
| Dussex et al. 2021 | Kākāpō | SLiM | mean $s = -0.024^*$ | Has purging occurred in the Stewart island kakapo population? |
| Grossen et al. 2020 | Alpine ibex | nemo | mean $s = -0.01^*$ | How has deleterious variation been impacted by a recent human-mediated bottleneck? |
| Kyriazis et al. 2022 | North American moose | SLiM | Kim et al. 2017<sup>^</sup> | How have bottlenecks influenced purging and genetic load in North American moose? |
| Marsden et al. 2016 | Dogs | PReFerSim | Boyko et al. 2008 | How has the domestication bottleneck shaped deleterious variation in dogs? |
| Robinson et al. 2018 | Channel island fox | SLiM | Kim et al. 2017 | How has recessive deleterious variation been impacted by small population size in island foxes? |
| Robinson et al. 2019 | Gray wolf | SLiM | Kim et al. 2017 | How does the large North American wolf population size influence recessive deleterious variation? |
| Robinson et al. 2022 | Vaquita | SLiM | *Estimated by authors*<sup>^</sup> | Are vaquitas doomed to extinction by inbreeding depression? |
| Stoffel et al. 2021b | Soay sheep | SLiM | Eyre-Walker et al. 2006<sup>#</sup> | Are short runs of homozygosity purged of deleterious variation? |
| Takou et al. 2021 | *Arabodopsis lyrata* | PReFerSim | *Estimated by authors* | Do range-edge populations have elevated genetic load? |
| Xie et al. 2022 | Chinese crocodile lizard | SLiM | Kim et al. 2017 | Have population declines resulted in purging? |

\*mean for gamma distribution, not based on explicit analysis
<sup>#</sup>DFE uses shape parameter from Eyre-Walker et al. 2006 and mean $s$ of -0.01, -0.03, -0.05
<sup>^</sup>sensitivity analysis conducted with additional DFEs

We suggest that future research should incorporate similar approaches to those implemented in these studies to provide a more thorough investigation of load in wild populations.

### What are the components of a simulation of deleterious genetic variation?

Modelling deleterious genetic variation in a simulation framework at a minimum requires specifying a population history, mutation rate, recombination rate, deleterious mutation target size, and distribution of selection and dominance coefficients (Table 2). The extent to which these parameters need to be tailored to a focal organism will vary depending on questions being asked. Many studies may be interested in using simulations primarily to explore qualitative dynamics of deleterious variation under various demographic and genetic scenarios. For example, one may be interested in asking: what are the qualitative effects of a bottleneck on genetic load under two extreme scenarios where deleterious mutations are either fully additive or fully recessive? For these studies, tailoring the simulation parameters to the focal organism may not be crucial, so long as the chosen parameters are reasonable.

**Table 2:**
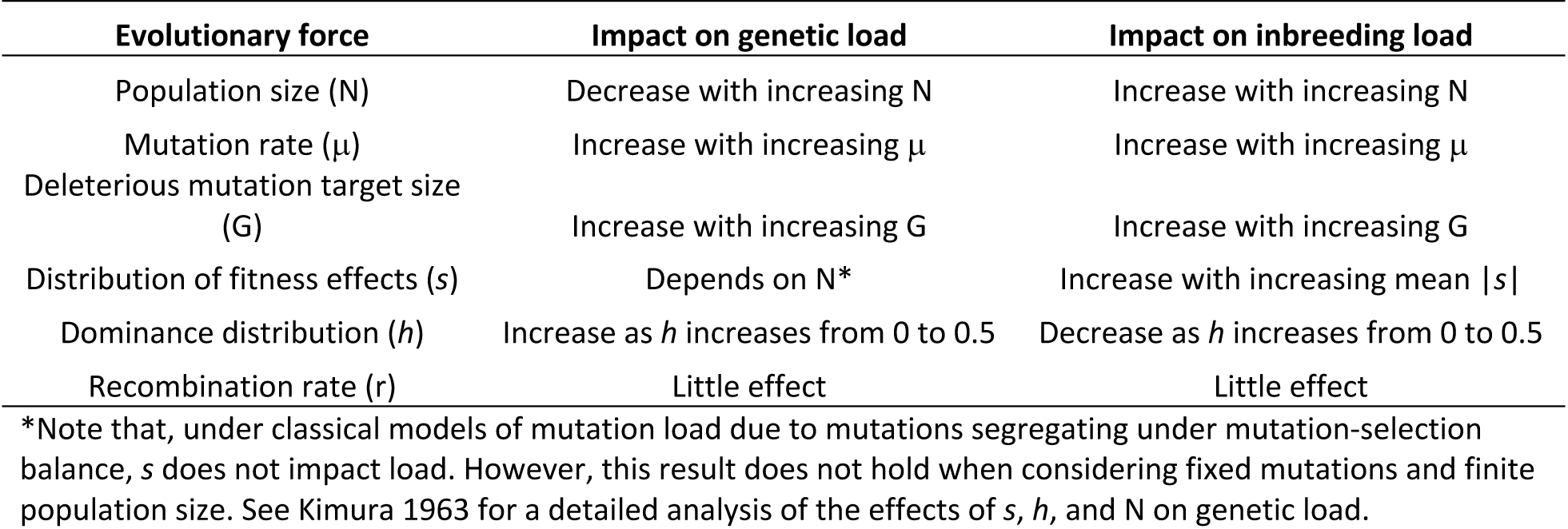
Evolutionary forces relevant to modelling load and how these forces impact load.

For studies aiming to make more quantitative statements about genetic load or project future extinction risk, tailoring simulation parameters to the focal organism may be more critical. For example, demographic history can vary widely between populations and has a large influence on deleterious genetic variation, and is therefore a crucial factor in modelling load and extinction risk. Fortunately, historical demographic parameters can be inferred from genomic datasets, though estimating recent demography (i.e., during the last tens or hundreds of generations) remains challenging (reviewed in (55)). The mutation rate is another essential component influencing levels of deleterious variation in a population, though high-quality mutation rate estimates (i.e., based on a large number of sequenced trios) do not exist for the vast majority of species. However, mutation rates can also be estimated from substitution rates between species, an approach that is now widely feasible given the abundance of whole genome sequencing data (56). Precisely estimating recombination rates may be somewhat less important for modelling load, though a growing number of approaches exist for estimating recombination rates from genomic datasets from as little as one diploid individual (e.g., (57, 58)). Tailoring the genome structure (i.e., the length and number of genes and extent of non-coding variation, which together determine the deleterious mutation target size) of a simulation to a specific organism can also be an important component of a simulation, particularly for studies aiming to model extinction risk in more ecologically-realistic models (51, 59). To aid in this, a growing number of annotated reference genomes are now available, which can provide useful information on genome structure, particularly for protein-coding regions of the genome (60).

Finally, the joint distributions of selection and dominance coefficients are essential components of modelling deleterious variation and load. These distributions determine the effect that new mutations exert on fitness, as well as the corresponding dominance coefficient of a mutation based on its selection coefficient. Although there is broad agreement that more deleterious mutations tend to be more recessive, the parameters of the distribution of dominance coefficients remain especially poorly known (61–64). Much more is known about the distribution of selection coefficients for new mutations, often termed “the distribution of fitness effects” or DFE, though most studies remain focused on humans and model organisms such as *Drosophila* ((65–67); Fig. 1). Given the importance of the DFE for simulations of deleterious variation, as well as recent debate over DFE parameters (68–70), below we provide a more detailed review of this topic.

**Figure 1:**
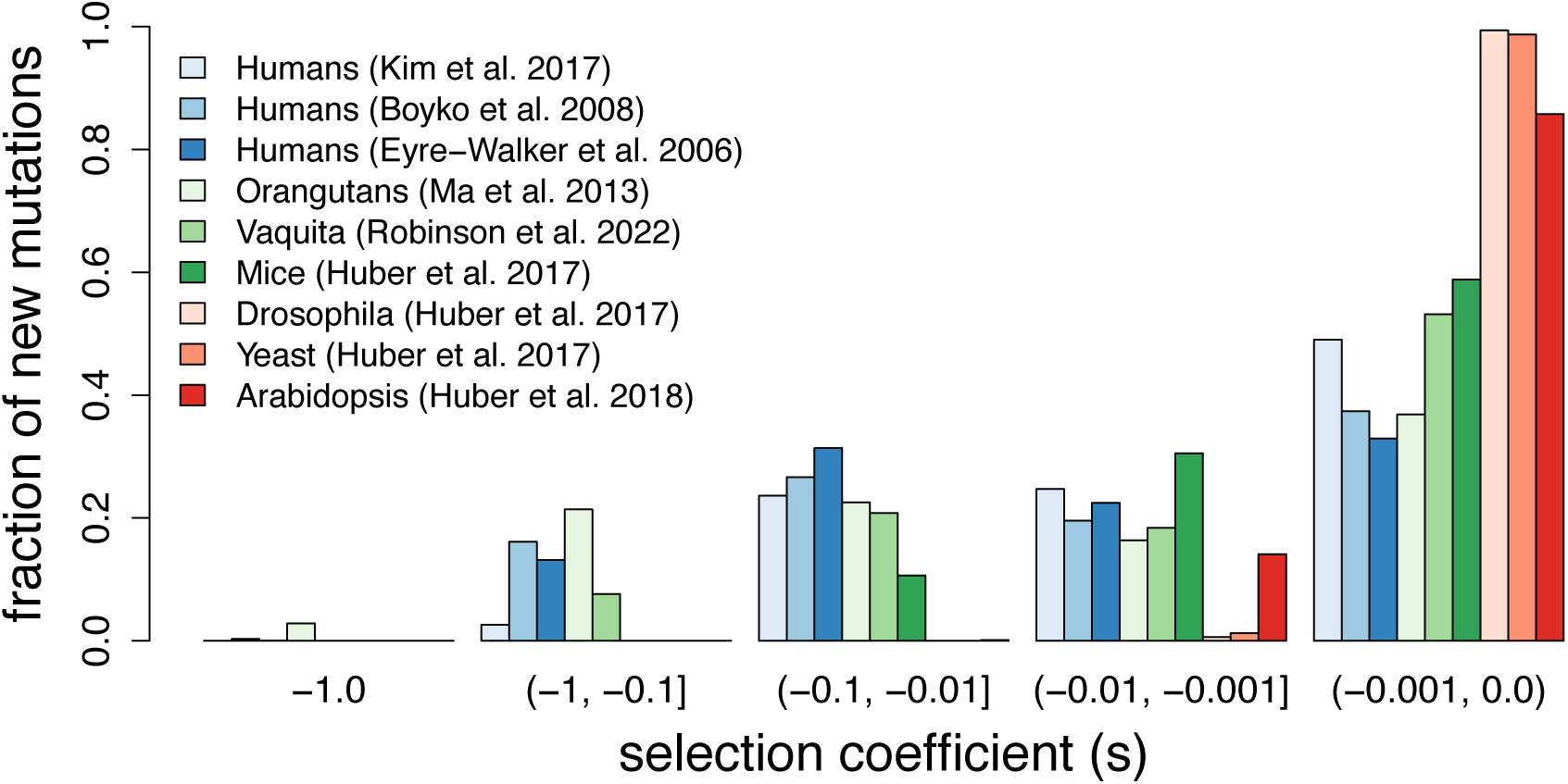
Representative estimates of the distribution of fitness effects from sequence-based approaches. Distributions are plotted in discrete bins of weakly deleterious (s >= -0.001), moderately deleterious (−0.001 > s >= -0.01), strongly deleterious (−0.01 > s >= -0.1), semi-lethal (−0.1 > s > -1) and lethal (s = -1.0) mutations. DFEs estimated for humans are colored in shades of blue, DFEs for non-human mammals are in shades of green, and non-mammalian DFEs are in shades of red. Note the higher fraction of weakly deleterious mutations in non-mammalian DFEs.

### What is the DFE and how is it estimated?

The DFE is a probability distribution that quantifies the selective effect (*s*) of new mutations entering the population, i.e., what fraction of new mutations are adaptive, neutral, weakly deleterious, or strongly deleterious. Here, we focus our discussion on the deleterious portion of the DFE, given that adaptive mutations do not influence load. Importantly, the DFE is not an estimate of segregating variation and therefore does not directly quantify load (see Supplemental Appendix 1; Fig. S1), a misconception that has recently spread in the literature (e.g., (71, 72)). Instead, the fate of a mutation after it enters a population, and whether it will segregate and potentially reach fixation, will be influenced by selection as well as the stochastic effects of genetic drift and linkage. Thus, quantifying segregating variation and load using the DFE requires modelling these effects under a given demographic model (see Supplemental Appendix 1 for an example; Fig. S1).

Historically, the DFE was estimated primarily using experimental mutation accumulation approaches (63, 65, 73, 74). However, these approaches are limited to detecting the small fraction of deleterious mutations that have large enough effects to be observed in a laboratory setting ((65, 74, 75); see Supplemental Appendix 2; Table S1). These limitations motivated the development of sequence-based approaches for estimating the DFE over the past two decades (65). Sequence-based methods estimate the DFE based on differences in the synonymous (assumed to be neutral) and nonsynonymous (assumed to be primarily neutral and deleterious) site frequency spectra (SFS), a summary of allele frequencies in a sample ((65, 67, 76–78); see Supplemental Appendix 2). Specifically, these methods typically use the synonymous SFS to control for neutral demographic effects and, conditioning on inferred demographic or nuisance parameters, then estimate the parameters of the distribution of *s* for new nonsynonymous mutations (most frequently, the mean and shape parameters of a gamma distribution). Thus, although these approaches have much greater power for estimating the weakly deleterious portion of the DFE, existing sequence-based DFEs are generally limited to nonsynonymous single nucleotide variants (though see (79)). Finally, one important limitation of sequence-based approaches is that they typically assume that all mutations have additive effects on fitness, given that information on the distribution of dominance coefficients is very limited (though see (61)). Consequently, sequence-based DFE approaches may not be well powered for estimating the relatively small portion of the DFE that is highly recessive and strongly deleterious (80).

### What are the best available DFE estimates?

A growing number of studies have used sequence-based methods to estimate the DFE for nonsynonymous mutations in various taxa including humans, non-human primates, mice, *Arabidopsis, Drosophila*, and the highly endangered vaquita porpoise ((51, 61, 66, 67, 76–78, 81–83); Fig. 1). In general, these studies estimate a relatively high proportion of weakly deleterious mutations (here defined as *s* > -1e-3), though this fraction varies among major taxonomic groups. For example, studies in mammals generally estimate ∼50% of mutations as weakly deleterious, whereas studies in *Arabidopsis, Drosophila*, and yeast suggest that >80% of new nonsynonymous mutations are weakly deleterious (Fig. 1). Comparative analyses of the DFE have suggested that such differences may be related to species complexity (66) as well as life history traits, such as selfing (81, 84).

Recently, these sequence-based DFE estimates have been criticized for not properly estimating the high fraction of strongly deleterious mutations estimated by experimental studies (68–70). Specifically, Kardos et al. (69) and Pérez-Pereira et al. (68) both argued that DFEs with a high fraction of strongly deleterious variation and dominance distributions that are far less recessive are better supported by experimental studies. The model proposed by Kardos et al. (69) assumes a mean *s* = -0.05 augmented with an additional 5% of mutations being recessive lethal, implying ∼67% of new mutations being strongly deleterious (here defined as *s* < -0.01), whereas the Pérez-Pereira et al. (68) model assumes a mean *s* = -0.2, implying ∼71% of new mutations are strongly deleterious (Fig. 2; Supplemental Appendix 3). By contrast, one of the more recent sequence-based DFE estimates for nonsynonymous mutations in humans estimated a mean *s* = -0.0131 with ∼26% of mutations being strongly deleterious (67), a result that is generally concordant with other estimates in humans and non-human mammals (Fig. 1). Moreover, these sequence-based estimates are also in agreement with a broad literature in population genetics and functional genomics suggesting that the majority of nonsynonymous mutations have relatively minimal effects on fitness (85–87).

**Figure 2:**
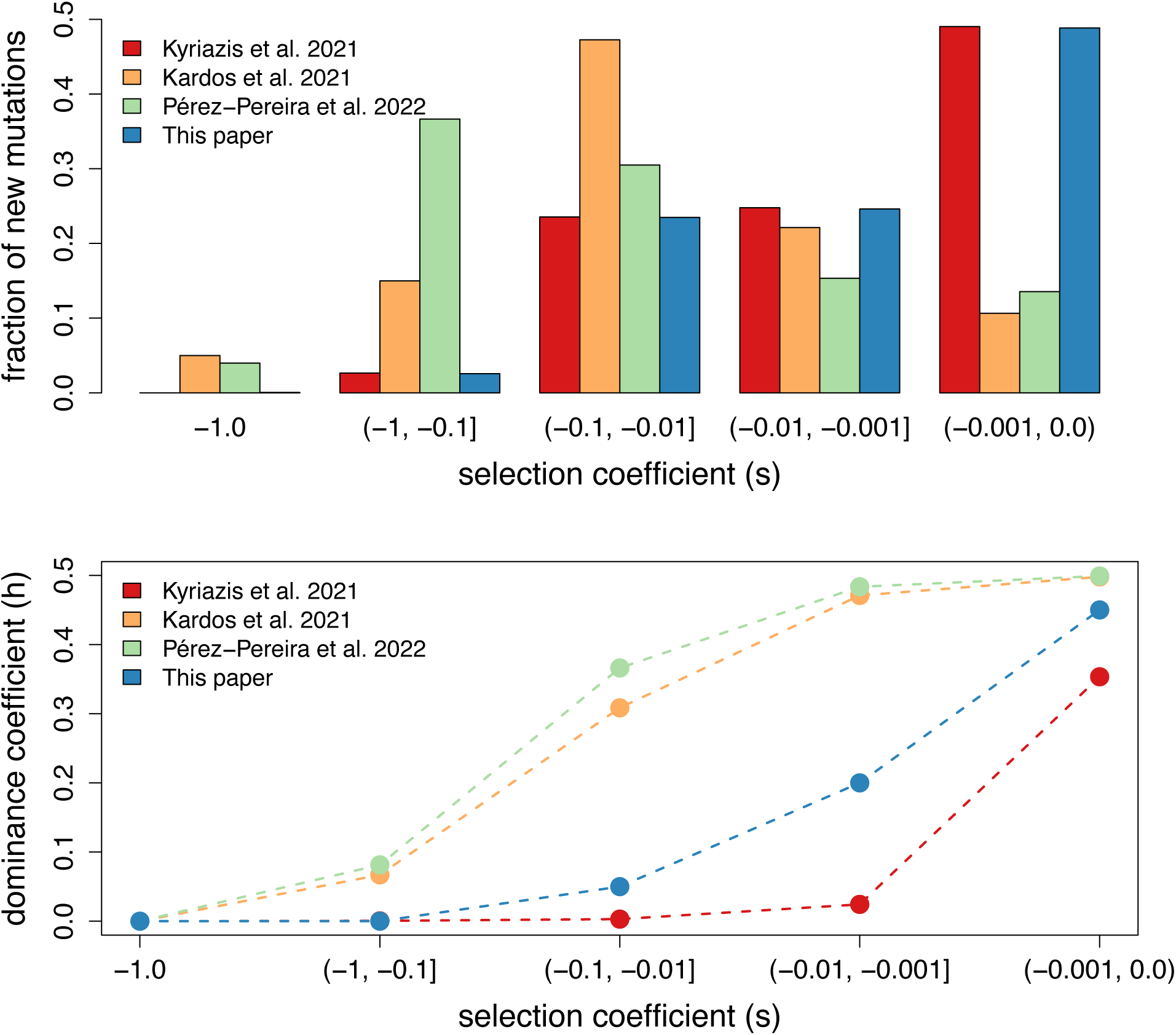
Comparison of DFE and dominance models employed by Kyriazis et al. 2021, Kardos et al. 2021, Pérez-Pereira et al. 2022, as well as the ‘best available’ model described in this paper. Distributions of *h* and *s* are plotted in discrete bins of weakly deleterious (*s* >= -0.001), moderately deleterious (−0.001 > *s* >= -0.01), strongly deleterious (−0.01 > *s* >= -0.1), semi-lethal (−0.1 > *s* > -1) and lethal (*s* < -0.99) mutations. Note that the ‘best available’ model includes 0.3% of mutations being recessive lethal based on results from Wade et al. 2022. Also note that the dominance distribution from Pérez-Pereira et al. 2022 assumes a distribution of *h* for each value of *s*.

The observation that sequence-based DFEs contain a much smaller proportion of highly deleterious mutations compared to experimental-based DFEs is not surprising, given that experimental studies are known to be biased towards detecting strongly deleterious mutations (65, 75) and sequence-based studies may be somewhat underpowered to detect such variation (80). However, if experimental-based studies are presumed to estimate the full DFE for nonsynonymous mutations, they should nevertheless make predictions that are consistent with patterns of nonsynonymous variation in genomic sequencing datasets. To investigate whether the Pérez-Pereira et al. (68) and Kardos et al. (69) models are supported by such information, we ran simulations under a human demographic model (67) where we compared the predicted nonsynonymous SFS from to the empirical SFS from the 1000G dataset for humans (88). We find that both models exhibit poor agreement with the empirical 1000G SFS, predicting nonsynonymous SFS that are greatly shifted towards rare mutations (Fig. 3; Table S2). For example, the Kardos et al. (69) and Pérez-Pereira et al. (68) models predict ∼72-76% of nonsynonymous mutations to be singletons (variants with frequency 1/2n), whereas ∼57% of variations are singletons in the 1000G dataset (Fig. 3; Table S2). This surplus of rare variation is due to the very strong predicted effects of negative selection under these models, which results in deleterious mutations being held at low frequency. This result is consistent with the expectation that such models based on experimental studies are biased towards highly deleterious mutations (65, 74, 75). By contrast, the Kyriazis et al. (59) model, which consists of the Kim et al. (67) DFE and a dominance distribution proposed by Henn et al. (25), is in much better agreement with empirical data, predicting ∼55% of variants as singletons (Fig. 3; Table S2). Thus, these results demonstrate that the Kardos et al. (69) and Pérez-Pereira et al. (68) models make predictions that are inconsistent with patterns of genetic variation in human genomic sequencing datasets (see Supplemental Appendix 4 for further discussion).

**Figure 3:**
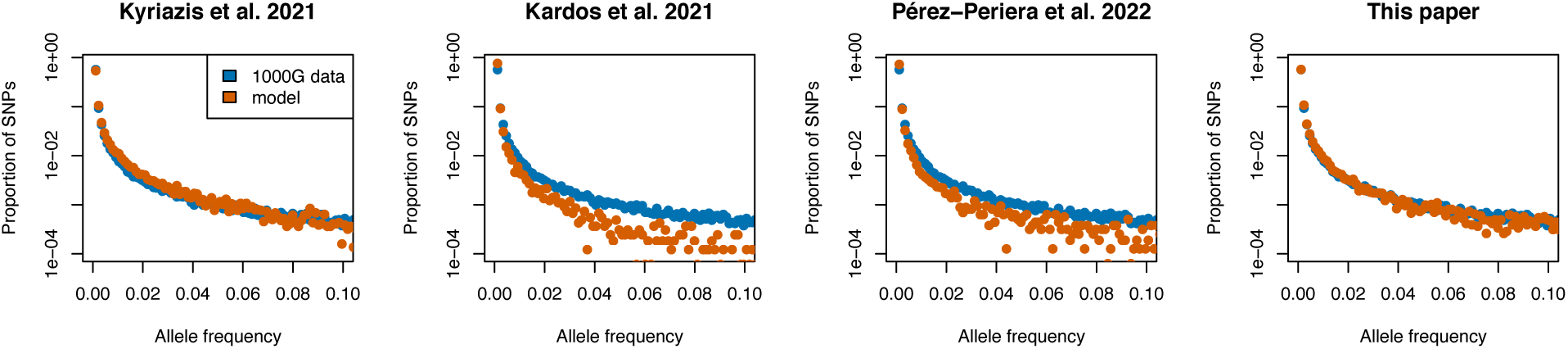
Predicted proportional nonsynonymous SFS from various DFE and dominance models compared to SFS from 1000G data. Predicted nonsynonymous SFS derived from simulations of ∼8.1 Mb of coding sequence under a human demographic model inferred using the synonymous SFS from the same 1000G dataset. Note that the predicted SFS from the Kyriazis et al. 2021 model and the ‘best available’ model proposed in this paper fit the 1000G data well, whereas the predicted SFS from the Pérez-Pereira et al. 2022 and Kardos et al. 2021 models are greatly shifted towards rare alleles due to the overabundance of strongly deleterious variation in these models. Also note that many entries of the Pérez-Pereira et al. 2022 and Kardos et al. 2021 SFS about allele frequency 5% are 0 and are therefore are omitted due to log scaling. See Figure S3 for plots of simulated vs empirical synonymous SFS and Table S2 for proportion of singletons and common variants predicted by each model.

Another way to validate proposed DFE and dominance models is to compare the predicted inbreeding load from each model to empirical estimates of inbreeding load (2B). Such empirical estimates can be derived by regressing measurements of fitness against estimates of individual inbreeding in a population (3, 4, 15, 89, 90). These estimates therefore represent an orthogonal source of information on deleterious mutation parameters. When comparing the predicted inbreeding load from the Kardos et al. (69), Pérez-Pereira et al. (68), and Kyriazis et al. (59) DFE and dominance models under simulations assuming human genomic and demographic parameters (i.e., assuming a genomic deleterious mutation rate of U=0.63; see Supplemental Appendix 3), we find that all models over-predict the estimated inbreeding load in humans of 2B=1.4 (91), as well as the median estimated inbreeding load in captive mammals of 2B=3.1 (90) and wild vertebrates of 2B=4.5 (89). Specifically, the Kardos et al. (69) and Pérez-Pereira et al. (68) models very high inbreeding loads of 2B=20.0 and 2B=28.4, respectively, while the Kyriazis et al. (59) model also predicts a substantial inbreeding load of 2B=11.3 (Fig. 4). Additionally, the partitioning of the inbreeding load (i.e., the contribution of the inbreeding load from detrimental mutations, semi-lethal mutations, and lethal mutations) in these models is also not consistent with empirical measures. For example, the Kardos et al. (69) and Pérez-Pereira et al. (68) models predict 16.0 and 12.2 recessive lethal mutations per individual, respectively (Fig. 4), whereas an empirical estimate suggests 0.6 recessive lethal mutations per human (92). By contrast, the Kyriazis et al. (59) predicts no recessive lethal mutations (see Supplemental Appendix 4 for further discussion). Thus, none of these models are consistent with empirical inbreeding load estimates, though over-predictions are especially notable for the Kardos et al. (69) and Pérez-Pereira et al. (68) models, due to these original analyses assuming unrealistically low effective population sizes (see Supplemental Appendix 5). Importantly, this over-prediction of the inbreeding load becomes much more extreme when assuming the original genomic deleterious mutation rate (U=1.2) of the Kardos et al. (69) model, and somewhat less extreme when assuming the original genomic deleterious mutation rate (U=0.4) from the Pérez-Pereira et al. (68) model (Fig. S5).

**Figure 4:**
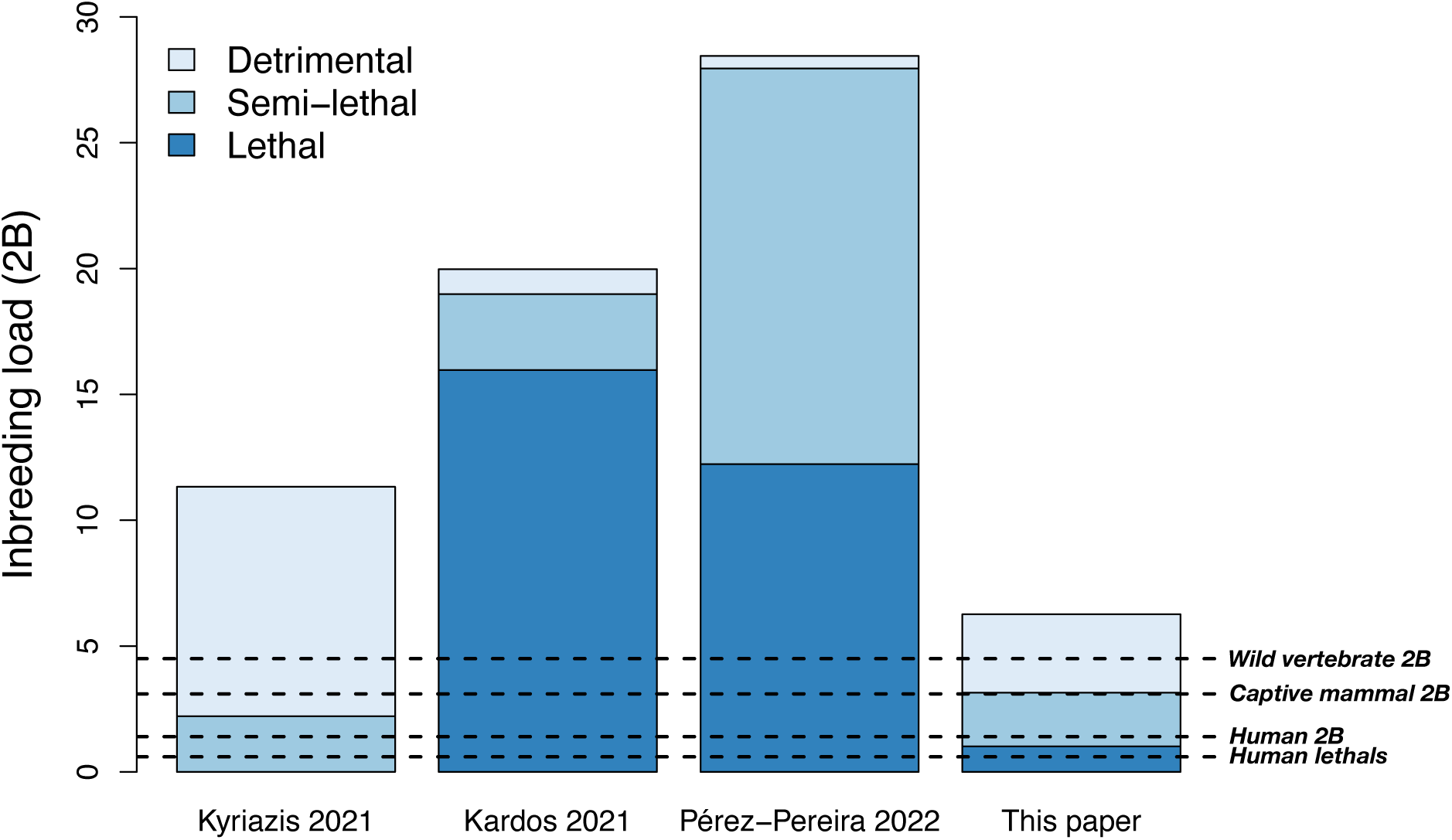
Inbreeding load partitioning under various DFE and dominance models using demographic and genomic parameters for humans. Colors depict contribution of inbreeding load from each class of deleterious mutations, with the total height of each bar representing the total predicted inbreeding load (2B). Detrimentals are here defined as mutations with *s* > -0.1, semi-lethals as mutations with -0.99 < *s* <= -0.1, and lethals as mutations with *s* <= -0.99. Dashed lines show estimated of number of lethals per diploid human from Gao et al. 2015 (“human lethals”), inbreeding load estimate for humans from Bittles & Neel 1994 (“human 2B”), and estimate of average inbreeding load for vertebrates from Nietlisbach et al. 2018 (“vertebrate 2B”). Note that predicted inbreeding load partitioning under the model proposed in this paper agrees well with empirical estimates, whereas predicted inbreeding load partitioning from other models do not. All displayed models assume a genomic deleterious mutation rate of U=0.63. See Fig. S5 for results under differing mutation rates.

Given the shortcomings of these existing models, we propose a new model based on an analysis of the DFE in humans under non-additive model (93) as well as an estimation of the recessive lethal portion of the DFE (80). In brief, this ‘best available’ model assumes a somewhat less recessive dominance distribution compared to the Henn et al. (25) distribution assumed by Kyriazis et al. (59) model and is augmented with a small proportion (0.3%) of recessive lethal mutations (see Supplemental Appendix 3 for details). Indeed, simulation results under this model are in much better agreement with empirical estimates. Specifically, our best available model predicts an inbreeding load of 2B=6.3, which is relatively evenly partitioned into contributions from detrimentals, semi-lethals, and lethals (Fig. 4), in agreement with empirical evidence (63, 92, 94, 95). Although this value exceeds empirical estimates in humans (2B=1.4) and captive mammals (2B=3.1), this result is expected given that these estimates are based on juvenile survival and may therefore be underestimates (90, 91). Overall, this analysis demonstrates that sequence-based DFE estimates can explain empirical patterns of the inbreeding load when making slight adjustments to account for their shortcomings in estimating the proportion of recessive lethal mutations (80). Thus, these DFEs remain preferable for modelling deleterious variation in coding regions in that they account for the impacts of both weakly and strongly deleterious variation. To facilitate use of our best available model in future simulation studies, we have provided an example SLiM script available on GitHub https://github.com/ckyriazis/simulations_review.

### Comparing genomics-informed models of inbreeding load to traditional PVA approaches

Although simulation-based approaches remain under-used in genomic studies of genetic load in wild populations, simulations have long been employed in conservation genetics in the context of population viability analyses (PVAs; (43–46)). For example, the simulation software VORTEX has been widely used to model inbreeding depression in wild populations (43, 96). Such programs allow the user to specify an empirical estimate of the inbreeding load for their focal organism. However, a key limitation of this approach is that empirical estimates of the inbreeding load do not exist for the vast majority of species, and many existing estimates are not reliable (89). Accurately estimating the inbreeding load is not a trivial task: it requires large sample sizes, accurate estimates of the inbreeding coefficient ideally from genomic data, high variance in inbreeding in a population, and a reliable proxy for fitness (89, 97). Relatively few studies exist that combine all of these elements, leading to wide variance in available estimates (see Supplemental Appendix 6 for further discussion).

Given the relative lack of empirical estimates of the inbreeding load, most existing PVA studies of inbreeding depression instead employ a default inbreeding load of 2B=3.1, a value derived from captive mammals by Ralls et al. (90). Many studies opt to use a much higher value of 2B=12 from O’Grady et al. (98), though this estimate has been shown to be unreliable ((89); see Supplemental Appendix 6). This use of a default, one-size-fits-all inbreeding load estimate is a critical limitation for many existing PVA studies, as it ignores the possibility of the inbreeding load varying across species and being influenced by the genetic and demographic characteristics of a given population. This issue may be most important for species with long-term small population size, for which inbreeding load is likely to be reduced (51, 59).

A genomics-informed simulation approach for modelling inbreeding depression offers many potential advantages to help overcome the limitations of using default inbreeding load values. In a genomics-informed approach, inbreeding load is modeled as an emergent property of fundamental genetic and demographic parameters that can be estimated from genomic data, rather than being predetermined. Based on these parameters, such models can predict the inbreeding load for a given species, obviating the need of a default estimate. The benefits of a genomics-based approach are illustrated by the analysis of extinction risk for the critically endangered vaquita porpoise presented in Robinson et al. (51), a species for which the inbreeding load is expected to be low due to long-term small population size (Fig. S4), though no empirical estimate of the inbreeding load exists for this population. By parameterizing simulations with genomics-based estimates of demographic parameters, mutation rates, and the DFE, we predicted an inbreeding load of 2B=0.95 and a high potential for recovery in the absence of excess mortality driven by the use of illegal gillnets. Moreover, we demonstrated that this low inbreeding load is largely a consequence of the small historical population size of the vaquita, finding that simulations with a 20x increased historical population size resulted in an increased predicted inbreeding load of 2B=3.32 and a much lower potential for recovery. Thus, the conclusion of high recovery potential for the vaquita would not have been reached when assuming a default inbreeding load for mammals of 2B=3.1.

There are several potential drawbacks to a genomics-informed simulation approach for modelling inbreeding depression, which should be taken into consideration before attempting to employ such an approach in a PVA model. First, forward-in-time simulations of genome-scale genetic variation tend to be highly computationally intensive, particularly when modelling large effective population sizes (N_e_ >> 10,000). This high computational load is largely due to the need for long burn-in periods for each simulation replicate during which mutations are allowed to reach their equilibrium frequency, a process that typically takes 10*N_e_ generations. However, these long burn-ins can be shortened by instead using durations long enough for inbreeding load to reach equilibrium, which typically takes <1*N_e_ generations (Fig. S6). Second, the reliability of a genomics-based PVA depends in large part on obtaining accurate estimates of relevant genetic and demographic parameters, which may be challenging for many species. In particular, mutation rates can be especially difficult to estimate and have a large influence on the predicted inbreeding load in a model. For example, Robinson et al. (51) estimated a mutation rate for the vaquita of 5.8e-9 per site per generation using a substitution-based approach, though with a plausible range of 2.2e-9 and 1.08e-8. When conducting sensitivity analyses under these varying mutation rates, we found that the predicted inbreeding load ranged from 2B=0.2 to 2B=2.4 and these varying inbreeding loads led to notable differences in projected extinction rates (51). Thus, genomics-based PVAs should be subject to extensive sensitivity analyses for genetic and demographic parameters and, given the challenges of validating these models, should be interpreted cautiously, as is the case for any PVA model (45).

Finally, although our discussion here focuses on modelling the influence of deleterious variation and genetic load on extinction, numerous other genetic threats to extinction can also be included in a genomics-informed PVA using modern simulation tools. For example, increased susceptibility to disease due to low genetic diversity could contribute to population declines (99), as can the challenge of adapting to changing environmental conditions (100). Future work should aim to better parameterize these processes in wild populations in order to more fully integrate evolutionary processes into PVAs (101).

### Remaining questions

#### How much does the DFE and partitioning of inbreeding load differ across taxa?

Much of our analysis in this paper focuses on the human DFE, given that genomic and demographic parameters in humans have been subject to extensive study. However, the extent to which the DFE may differ across species remains poorly known. Although available DFE estimates in mammals are generally similar to humans (Fig. 1), relatively few species have been examined, and little to no information is available on the DFE in non-mammalian vertebrates. Obtaining a better understanding of the DFE across diverse vertebrate species will help determine whether it is justified to use mammalian DFEs, such as the human DFE presented in this paper (Fig. 2), in simulations for other vertebrate taxa where DFE estimates are not available. Moreover, additional DFE estimates will also be informative as to the extent to which inbreeding load partitioning may vary across taxa.

#### What is the distribution of dominance coefficients?

Minimal information exists on the distribution of dominance coefficients, and the few studies that exist are based on a handful of species (61–64). Moreover, results from these studies sometimes conflict with one another, with available evidence in yeast and *Drosophila* suggesting a much less recessive distribution of dominance coefficients compared to results from *Arabidopsis* (61–64). Whether this discrepancy is a consequence of true biological differences among these taxa or is instead due to methodological issues remains unclear.

Obtaining additional information on the distribution of dominance coefficients represents one of the most essential components for studies modelling inbreeding depression. Specifically, if most deleterious mutations are found to have dominance coefficients that are highly recessive (*h*<0.05) as suggested by a recent study (61), this would suggest a high severity of inbreeding depression due to contributions from both weakly and strongly deleterious mutations. Moreover, a highly recessive dominance distribution would also suggest a very large dependence of inbreeding depression on historical population size (59, 102, 103).

#### What is the role of heterozygote advantage and non-coding variation in inbreeding depression?

Understanding the genetic basis of inbreeding depression has long been a topic of interest in evolutionary genetics (104). Although many of the simulation models discussed above focus solely on the contribution of recessive deleterious mutations at nonsynonymous sites in coding regions (e.g., (51, 54, 59)), it is likely that other mechanisms and types of mutations also contribute to some extent. For example, heterozygote advantage could play a role in influencing inbreeding depression, though its contribution has historically been hard to quantify (104). Similarly, deleterious mutations in conserved noncoding regions could also contribute, given that ∼4.5% of the non-coding genome has been shown to be highly conserved across mammals, potentially implying a high strength of negative selection in these regions (10, 22, 105, 106).

Although these mechanisms and types of mutations likely play some role in influencing inbreeding depression, available evidence suggests that their impact may be relatively minimal. Specifically, studies on heterozygote advantage generally suggest that it occurs only at a small number of loci in the genome (107), implying a small contribution to inbreeding depression (104). Similarly, although deleterious mutations in conserved noncoding regions appear to be widespread, several studies suggest that these mutations tend to be weakly deleterious (*s* on the order of 1e-3; (79, 108, 109)), such that they likely do not contribute much to inbreeding depression. These considerations imply that recessive deleterious mutations in coding regions likely represent the predominant driver of inbreeding depression. In support of this, we note that the models of inbreeding depression summarized in Figure 4 all ignore contributions of these mechanisms, yet all of these models predict inbreeding loads that are at least as large as those observed empirically. Nevertheless, we emphasize that additional research is needed to explore the potential contribution of heterozygote advantage and noncoding variation to inbreeding depression and extinction risk. For example, simulation studies that parameterize these factors based on the literature and quantify their effect on inbreeding depression would be a valuable step towards better determining their importance.

#### How do simulation dynamics in a Wright-Fisher model compare to those in more ecologically-complex models?

The Wright-Fisher (WF) model is a ubiquitous model in population genetics that underlies many of the methodological approaches discussed above. For example, sequence-based DFE methods assume a WF model (67, 76–78, 110), as do the simulation results presented in Figures 3 and 4. However, in many cases, it may be desirable to use parameter estimates derived from a Wright-Fisher model in the context of more ecologically complex simulation models, such as the non-Wright-Fisher (nonWF) model in SLiM3 (36). This model allows for populations with overlapping generations, an important departure from the WF model that is also present in other ecologically-realistic simulation programs such as Nemo-age (39).

A key question when using these more complex models with overlapping generations is whether parameters that are inferred assuming a WF model apply in a nonWF setting. For example, DFEs inferred assuming a WF model are scaled in terms of per-generation selection coefficients rather than per-year. As models with overlapping generations assume selection occurs on a yearly timescale (or some other arbitrary unit of time), the selection coefficients from the DFE will need to be rescaled. Although theory suggests that selection coefficients in this case can be rescaled simply by dividing by the generation time of a species (111), the extent to which this rescaling should apply to the full DFE is not clear. For example, highly deleterious mutations that impair development are likely to act during early life stages and not over the span of an entire generation, thus dividing the selection coefficient of such mutations by the generation time may have unintended consequences. In Robinson et al. (51), our solution to this issue was to rescale selection coefficients for weakly deleterious mutations with *s* > -0.01, but not for strongly deleterious mutations with *s* < -0.01. The validity of this approach, and more broadly the best practices for using parameter estimates derived from WF models under more complex models, remains in need of further investigation.

## Acknowledgements

We are grateful to Phil Morin, Annabel Beichman, Robert Wayne, Stella Yuan, and Meixi Lin for comments on the manuscript. C.C.K. and K.E.L. were supported by National Institutes of Health grant (R35GM119856 to K.E.L.).

## Data Availability

All simulation and plotting scripts are available at https://github.com/ckyriazis/simulations_review

## Author Contributions

C.C.K., J.A.R., and K.E.L. conceived the study. C.C.K. conducted analyses and wrote the manuscript with input from all authors.

