## Supplementary Appendix for "Using computational simulations to quantify genetic load and predict extinction risk"

**Table of Contents**

1. Using simulations to model segregating variation and estimate load
2. Overview of approaches for DFE inference
3. Description of simulation models
4. Validation of simulation models
5. The influence of effective population size on inbreeding load
6. Discussion of empirical inbreeding load estimates
7. Supplementary Figures and Tables

**1. Using simulations to model segregating variation and estimate load**

Here, we describe an example simulation to aid in demonstrating the differences between the distribution of *s* for new, segregating, and fixed mutations as well as illustrate how simulations can be used to quantify load. For this simulation, we used our 'best available' model with human genomic parameters as described below. We modelled three effective population sizes of 50, 500, and 5000. For each simulated population, we ran a burn in for a duration of $10*N_e + 1000$ generations (i.e., a burn in of 1500 generations for $N_e$=50 and 5100 generations for $N_e$=5000). These additional 1000 generations were included to allow each simulated population to accumulate fixed mutations following the initial burn-in for a period of time that was independent of $N_e$. We averaged across 10 simulation replicates for all results described below.

From the SLiM simulation output, it is possible to directly obtain the values of the selection coefficients (*s*) for segregating and fixed mutations. This enables comparison of the distribution of *s* for new, segregating, and fixed mutations. When comparing these distributions of *s* for new, segregating, and fixed mutations (Fig. S1), several key conclusions are evident. First, these distributions greatly differ, and the extent to which they differ depends on $N_e$. Specifically, weakly deleterious mutations ($s > -0.001$) comprise a larger fraction of segregating variation compared to the rate at which they enter the population, whereas more strongly deleterious mutations with $s < -0.001$ comprise a lower proportion of segregating variants compared to the rate at which they enter the population (Fig. S1). These effects are even more pronounced for fixed mutations, which tend to be largely weakly deleterious, though the extent to which this is true depends greatly on $N_e$. For example, all fixed mutations are weakly deleterious when $N_e$=5000, whereas a notable proportion of fixed mutations are moderately ($-0.01 < s =< -0.001$) or strongly deleterious ($-0.1 < s =< -0.01$) when $N_e$=50 (Fig. S1). These dynamics are reflective of role of $N_e$ in determining the effectiveness of negative selection against deleterious alleles: negative selection serves to remove deleterious mutations from the population, but the extent to which this occurs can be impaired by increased genetic drift at low $N_e$. Thus, the patterns of deleterious variation segregating or fixed in the population depend on both the initial DFE for new mutations as well as the demographic history of the population.

These considerations also emphasize that the DFE alone is not sufficient information to quantify genetic load. In a simulation, however, genetic load and inbreeding load can be easily quantified given that information of segregating and fixed deleterious mutations is readily available. To illustrate this, we estimated genetic load and inbreeding load from these simulations using the equations described in the main text. For example, for two loci with $q_1$=0.01, $s_1$=0.03, $h_1$=0.05 and $q_2$=0.3, $s_2$=0.001, and $h_2$=0.2, the mean genetic load (L) can be calculated using equation (1) from the main text as:

$$L = 1-\bar{w}$$

$$\bar{w} = \prod_i [1 - 2h_i s_i q_i(1 - q_i) - s_i {q_i}^2]$$

$$= [1-2h_1 s_1 q_1(1-q_1) - s_1 {q_1}^2]*[1-2h_2 s_2 q_2(1-q_2) - s_2 {q_2}^2]$$

$$= [1-2*0.05*0.03*0.01*0.99-(0.03*0.01^2)]*[1-2*0.001*0.2*0.3*0.7-(0.001*0.3^2)]$$

$$=0.9997933$$

$$L = 1-0.9997933 = 0.0002067$$

Similarly, the mean diploid inbreeding load (2B) can be calculated using equation (2) from the main text as:

$$2B = \sum_i [s_i q_i - s_i {q_i}^2 - h_i s_i q_i(1 - q_i)]$$

$$= 2[s_1 q_1 - s_1 {q_1}^2 - 2h_1 s_1 q_1(1-q_1)] + 2[s_2 q_2 - s_2 {q_2}^2 - 2h_2 s_2 q_2(1-q_2)]$$

$$= 2[0.03*0.01-0.03*0.01^2-2*0.05*0.03*0.01*0.99] +$$

$$2[0.001*0.3-0.001*0.3^2-2*0.2*0.001*0.3*0.7]$$

$$=0.000786$$

Note that, for these equations, selection coefficients for deleterious mutations are positive, though elsewhere we assume negative *s* for deleterious mutations.

Using this same approach applied to our simulation output, we estimate a mean genetic load of 0.69 when $N_e$=50, 0.42 when $N_e$=500, and 0.41 when $N_e$=5000, and a mean inbreeding load of 0.56 when $N_e$=50, 1.66 when $N_e$=500, and 3.84 when $N_e$=5000 (Fig. S2). Thus, genetic load and inbreeding load can greatly differ with the same underlying DFE due to the effects of demography. Moreover, changes in load with $N_e$ are not necessarily linear, as evidenced by the highly similar genetic loads of the $N_e$=500 and $N_e$=5000 populations. Lastly, how the genetic load correlates with $N_e$ may be in the opposite direction of how the inbreeding load correlates with the DFE. Overall, this analysis demonstrates that load due to segregating and fixed mutations can be estimated from a simulation, though doing so requires careful modelling of demography and the DFE.

### 2. Overview of approaches for DFE inference

Estimation of the distribution of fitness effects of new mutations represents a long-standing challenge in population genetics, employing a variety of different approaches. Historically, work focused on co-estimation of mean selection coefficient (E(*s*)) and genome-wide deleterious mutation rate (U) using experimental mutation accumulation (MA) studies (1–4). Results from this approach are varied, with some studies estimating a very low U on the order of 0.05 or lower and relatively large E(*s*) on the order of -0.05 or greater, whereas other studies estimate a much larger U on the order of 1 and smaller E(*s*) on the order of -0.001 (2).

Experimental approaches have numerous limitations, which may hinder their ability to accurately estimate the DFE. First, nearly all experimental studies suffer from the limitation that U and E(*s*) are confounded (2, 4, 5). In other words, the distributions of fitness in the laboratory populations can be explained by a high U and a low E(*s*), or vice versa. Second, most experimental studies also have wide confidence intervals in their estimates, given that very few mutations are typically observed, and results can be highly dependent on the analysis approach (2, 4, 5). Third, experimental studies can only detect mutations with very large effects on fitness, as the impacts of smaller-effect mutations cannot be observed in a laboratory setting (2, 4, 6). For example, Davies et al. (6) found that only 4% of new deleterious mutations in *C. elegans* can be detected using an experimental approach (6). Finally, experimental studies can only be conducted on organisms that are amenable to a laboratory setting, with most studies being conducted on *Drosophila*, yeast, *C. elegans,* and *Daphnia* (2, 5). Thus, the extent to which results from these studies can be extrapolated to other taxonomic groups remains in question.

To overcome these limitations, there has been a growing use of DNA-sequencing-based methods for estimating the DFE over the last two decades. These approaches typically focus on estimating parameters of DFE based on differences in synonymous (which is assumed to be neutral) and nonsynonymous (assumed to be primarily neutral and deleterious) site frequency spectra (SFS), a summary of allele frequencies in a sample (4, 7–10). Specifically, sequence-based approaches rely on the observation that genetic variation is typically depleted in the nonsynonymous SFS relative to the synonymous SFS, and this depletion is a consequence of negative natural selection against deleterious nonsynonymous alleles. This depletion can therefore be leveraged to estimate the parameters of the DFE (most frequently, the mean and shape parameters of a gamma distribution) under an evolutionary model that controls for effects of demography. Thus, estimates of the DFE from sequence-based approaches are generally only available for nonsynonymous mutations (though see (11)).

Sequence-based approaches to DFE estimation also suffer from several limitations. First, the assumption that synonymous mutations can serve as a proxy for neutral mutations may be violated in some cases. For example, studies in *Drosophila* and yeast have uncovered signatures of relatively strong selection at synonymous sites (12, 13), and the impacts of these factors on DFE inference have yet to be fully explored (though see (14)). Second, these methods also generally assume that all mutations have additive effects on fitness, given the limited information on the distribution of dominance coefficients (though see (15)). Finally, sequence-based DFE approaches may also be limited in their ability to disentangle the strongly deleterious portion of the DFE ($s$ on the order of -0.01) from the lethal and semi-lethal portions of the DFE ($s < -0.1$). This is because sequence-based DFE approaches infer the presence of strongly deleterious new mutations based on a lack of genetic variation in the nonsynonymous SFS relative to that expected under an evolutionary model. However, strongly deleterious mutations with even relatively small $s$ (on the order of -0.01) often may not segregate at appreciable frequency in small to modest sample sizes and may therefore leave similar patterns to lethal mutations in genetic variation datasets. In Table S1, we summarize different approaches for detecting deleterious mutations, which mutation types they are best suited for detecting, and the mean strength of selection that is roughly expected for each class of mutations.

### 3. Description of simulation models

Our analysis compares four DFE and dominance models using simulations. Each of these models assume that the DFE is gamma distributed (some including a point mass for lethal mutations) and that dominance coefficients are inversely related to selection coefficients. Below, we describe each of these models of the DFE, reporting E($s$) and shape parameter for each gamma DFE, the function used to specify dominance, and the deleterious mutation rate originally assumed for each model. Note that selection here is parameterized such that the heterozygote has fitness 1+$sh$ and homozygous derived has fitness 1+$s$, whereas many DFE inference approaches assume a heterozygote fitness of 1+2$sh$ and homozygote derived fitness of 1+2$s$ (e.g., (7, 8)).

1. **Kyriazis et al. 2021 model:** This model assumes a DFE inferred by Kim et al. (7) based on human genetic variation data and a dominance distribution proposed by Henn et al. (16). The DFE assumes E($s$)=-0.0131 and shape=0.186, and the dominance coefficient for mutations follows the relationship $h$=(0.5)/(1 - 7071.07*$s$). However, note that for the sake of computational efficiency, this model was simplified to a discrete dominance distribution where $h$=0.25 when $s$>-0.01 and $h$=0.0 when $s$=<-0.01 for the majority of simulations in Kyriazis et al. (17). Also note that although the Henn et al. (16) dominance model was not estimated based on data, the high degree of recessivity it implies is supported by a recent analysis of dominance in *Arabidopsis* (15). Finally, this model also assumed a deleterious mutation rate per diploid of U=0.42, roughly based on coding sequence parameters for dogs (18).

2. **Kardos et al. 2021 model:** This model assumes a DFE with E($s$)=-0.05 and shape=0.5, augmented with an additional 5% of mutations being recessive lethal and a dominance

model with $h=0.5*\exp(-13s)$. These DFE parameters are loosely based on results from mutation-accumulation experiments (4), and the dominance model is similarly based on an estimate of a mean $h$=0.36 from mutation-accumulation studies (19, 20). This model assumes a deleterious mutation rate per diploid of U=1.2, which is derived from a genome-wide analysis in *Drosophila* (21).

3. **Pérez-Pereira et al. 2022 model:** This model assumes a DFE with E($s$)=-0.2 and shape=0.33 and a dominance model where $h$ was drawn from a uniform between 0 and exp(7.6$s$), where 7.6 was chosen to result in mean $h$=0.283. These parameters appear to be largely informed by an analysis of purging of highly deleterious alleles in experimental *Drosophila* populations from Pérez-Pereira et al. (22), which concluded that a model with very high E($s$)=-0.3 and low diploid U=0.04 were consistent with observed changes in fitness. However, note that the model in Pérez-Pereira et al. (23) assumes a much larger U=0.4. Finally, note that our implementation of the dominance function in this model assumes a fixed $h=0.5*\exp(7.6s)$, given that it is not possible to draw $h$ from a uniform distribution in SLiM.

4. **Our 'best available' model:** The model we propose in this paper is similar to the Kyriazis et al. (17) model above, though with a few important changes. First, we assume an increased deleterious mutation rate per diploid of U=0.63, informed by available estimates of human coding sequence parameters (see discussion below). Next, although we assume the same E($s$)=-0.0131 and shape=0.186, we augment this with an additional 0.3% of recessive lethals, based on the analysis of (24). This results in a predicted number of recessive lethals per diploid of ~1 when simulating under human demography (Fig. 4), in good agreement with empirical estimates (25). Finally, this model also assumes a more intermediate degree of recessiveness compared to the Kyriazis et al. (17) model, informed by the analysis of Cavassim & Lohmueller (26). Specifically, we enforce a discrete dominance distribution with $h$=0.45 for weakly deleterious mutations ($s > -0.001$), $h$=0.2 for moderately deleterious mutations ($-0.01<s$

=< -0.001), *h*=0.05 for strongly deleterious mutations ($-0.1<s=<-0.01$), and *h*=0.0 for lethal and semi-lethal mutations ($s<-0.1$; Fig. 2).

**4. Validation of simulation models**

As discussed in the main text, Kardos et al. (27) and Pérez-Pereira et al. (23) critique sequence-based DFEs for underestimating strongly deleterious variation. Kardos et al. (27) make this argument based on the observation that many experimental estimates of the DFE suggest a relatively large E(*s*) on the order of -0.05. However, as noted above, experimental studies are biased towards strongly deleterious variation and are unable to tease apart the confounded effects of E(*s*) and the genomic deleterious mutation rate U. Specifically, results from experimental studies are generally consistent either with a large E(*s*) on the order of -0.05 and small U on the order of 0.05, or a small E(*s*) on the order of -0.001 and large U on the order of 1 (2). Despite these observations, Kardos et al. (27) couple their proposed large E(*s*) of -0.05 with a large U of 1.2 estimated in *Drosophila* (21). By doing so, Kardos et al. (27) are in essence extrapolating a large E(*s*) estimate that likely applies only to a small fraction of deleterious mutations to model deleterious variation at a genome-wide scale. One consequence of this assumption is that their model implies very strong selection on non-coding variation, whereas available evidence suggests that selection even in highly conserved non-coding regions is primarily weak (mean *s* on the order of -1e-3; (11, 28, 29)).

Pérez-Pereira et al. (23) use a similar approach of using experimentally-derived strong selection parameters to model deleterious variation at a much broader scale. Specifically, their proposed E(*s*)=-0.2 is informed by an analysis of purging of highly deleterious alleles in experimental *Drosophila* populations, which concluded that a model with very high E(*s*)=-0.3 and low diploid U=0.04 were consistent with observed changes in fitness (22). In this analysis, Pérez-Pereira et al. (22) acknowledge that this large E(*s*) estimate applies only to a small proportion of deleterious mutations, and that most deleterious mutations have much smaller effect that are hard to detect in an experimental setting. However, despite this acknowledgement, the analysis of Pérez-Pereira et al. (23) nevertheless applies a large E(*s*)=-0.2 to a much larger set of deleterious mutations (U=0.4). As in Kardos et al. (27), this application of extremely strong selection parameters at a genome-wide scale results in a model that is not consistent with a broad literature in population genetics and functional genomics suggesting that most deleterious mutations have relatively minimal effects on fitness (30–32).

Here, we discuss two approaches for validating the models Kardos et al. (27) and Pérez-Pereira et al. (23) to more concretely demonstrate the consequences of this use of experimentally-derived large E(*s*) estimates to model a much broader set of deleterious mutations. First, we wanted to assess whether these models can match patterns of genetic variation in empirical data. Even though Kardos et al. (27) and Pérez-Pereira et al. (23) did not estimate the parameters of their DFE from genetic variation data, if their models are a reasonable approximation of biology, they should still match patterns of varation in genomic datasets. To test this, we compare the predicted nonsynonymous site frequency spectra (SFS) from the above-described models to the SFS from 432 unrelated European individuals from the 1000G dataset (33). We ran simulations assuming human genomic and demographic parameters, including a mutation rate per site of $\mu$=1.5e-8 (34), 1023 genes per autosome (reflecting 22,500 total genes across 22 autosomes) each with a length of 1340 bp (35), and a ratio of nonsynonymous to synonymous mutations of 2.31:1 (36). For this analysis, we assumed the demographic parameters inferred by Kim et al. (7) based on the synonymous SFS from 432 European individuals from the same 1000G dataset (33). This demographic model estimated (moving forward in time) an ancestral population size of $N_a$=12,378 diploids, followed by a bottleneck 1048 diploids for 248 generations, population growth to 13,625 for 1,744 generations, and finally exponential growth for the final 497 generations to a current effective population size of 659,551 diploids (see Tables S1 and S2 in Kim et al. (7)). We ran burn-ins for 10*Ne generations at $N_a$=12,378 and, to facilitate computational feasibility, simulated genetic variation for only six chromosomes (~8.1 Mb of coding sequence).

As summarized in the main text, these simulations demonstrate that the Kardos et al. (27) and Pérez-Pereira et al. (23) models predict nonsynonymous SFS that greatly differ from the

1000G SFS. Specifically, the SFS predicted by these models are greatly shifted towards rare alleles relative to the 1000G SFS, reflective of the very strong selection implied by these models (Fig. 3). For example, the Kardos et al. (27) and Pérez-Pereira et al. (23) models predict that ~72-76% of nonsynonymous variants are singletons, whereas ~57% of variants are singletons in the 1000G dataset (Fig. 3; Table S2). Moreover, common variants with allele frequency >5% comprise only ~3-6% of SNPs in the SFS from these models, whereas common variants make up 13.3% of the 1000G SFS (Fig. 3; Table S2). By contrast, the predicted SFS from the Kyriazis et al. (17) model is in much better agreement with the 1000G SFS, predicting ~55% of variants as singletons and 10.4% of variants as being common (Fig. 3; Table S2). The concordance between the SFS predicted by the Kyriazis model and the empirical SFS is notable given that the Kim et al. (7) DFE assumed by Kyriazis et al. (17) was inferred assuming additivity but was modelled in this paper using a highly recessive distribution of dominance coefficients (Fig. 2). Finally, the 'best available' model we propose in this paper is also in good agreement with the 1000G SFS, predicting ~57% of variants as singletons and ~10% as common (Fig. 3; Table S2). Thus, these results demonstrate that the selection and dominance parameters of Kardos et al. (27) and Pérez-Pereira et al. (23) are not consistent with patterns of genetic variation in humans. Moving forward, we recommend that when new models of evolutionary genetic parameters are proposed, researchers should test to see whether they can predict basic summaries of genetic variation as a means to validate their models.

A second orthogonal approach to validate these DFE and dominance models is to compare predicted inbreeding load to empirical estimates of the inbreeding load. Although Kardos et al. (27) and Pérez-Pereira et al. (23) make these comparisons, they assume small effective population sizes relative to those observed in natural populations. This is a crucial oversight for their analysis, given that all models predict that inbreeding load depends greatly on $N_e$ (Fig. S6). For example, Kardos et al. (27) examined equilibrium $N_e$ ranging from 25 to 1500, finding that $N_e$=800 predicts the median inbreeding load estimate from captive mammals of 2B=3.1 ((37); see Fig. S3 in Kardos et al. (27)). However, estimates of equilibrium $N_e$ based on genome-wide diversity in mammals suggests median $N_e$ of ~22,000 (Fig. S4; see below for further discussion).

Notably, the $N_e$ range examined by Kardos et al. (27) is almost entirely within the 1% quantile of the observed distribution, suggesting that it is highly unrepresentative for mammals. This implies that the Kardos et al. (27) DFE, when modelled using realistic values of $N_e$ for mammals, should predict an inbreeding load that is substantially higher than that observed in natural populations. Although the analysis of Pérez-Pereira et al. (23) assumes a larger $N_e$=10,000, this value is still relatively low for mammals, and is many orders of magnitude lower than that of *Drosophila* ($N_e$ range ~1e5-5e6; (36, 38, 39)), which serves as foundation for much of their analysis (22).

To explore inbreeding load predictions from the Kardos et al. (27) and Pérez-Pereira et al. (23) models as well as the Kyriazis et al. (17) model under more realistic demographic parameters, we used the same simulation framework as described above based on parameter estimates for humans. Here, however, we simulated variation across 22 autosomes, yielding a genomic deleterious mutation rate of U=0.63 per diploid. We assume this same genomic deleterious mutation rate for all models to facilitate direct comparison of model predictions under human genomic parameters, though we also present results under the rates originally assumed by Kyriazis et al. (17), Kardos et al. (27) and Pérez-Pereira et al. (23) in Figure S5. Note that, to facilitate computational efficiency, we did not model neutral mutations for the inbreeding load partitioning results (Figs 4 and S5) and ran burn-ins for only 10,000 generations, which was long enough for the simulated inbreeding load to reach equilibrium (Fig. S6). To visualize results, we plot inbreeding load partitioned into three categories: load due to lethal mutations ($s \leq -0.99$), load due to semi-lethal mutations ($-0.99 < s \leq -0.1$) and load due to detrimentals ($s > -0.1$). We compare these predictions to the inbreeding load estimate in humans of 2B=1.4 (40), the segregating recessive lethal estimate in humans of 0.6 per diploid (25), the median inbreeding load estimate in captive mammals of 2B=3.1 (37), and the median inbreeding load estimate in wild vertebrates of 2B=4.5 (41). We do not include the estimate of 2B=12 for wild vertebrates from O'Grady et al. (42), as it has previously been shown to be unreliable ((41); see below for further discussion). Finally, we note that, although Nietlisbach et al. (41) report a mean 2B=7, we computed a median from their data to be consistent with the use of a median from Ralls et al. (37).

As summarized in the main text, these results demonstrate that all existing models greatly over-predict the empirical inbreeding load in humans and other species. For example, the Kyriazis et al. (17) model predicts a total inbreeding load of 2B=11.3, largely due to detrimentals. The lack of lethals in this model is consistent with the critique that sequence-based DFEs are unable to detect the presence of recessive lethals (24). However, this slight underestimation does not imply that the large fraction of recessive lethals assumed by Kardos et al. (27) and Pérez-Pereira et al. (23) is correct. Specifically, the models of Kardos et al. (27) and Pérez-Pereira et al. (23) greatly overshoot the empirical estimate of ~0.6 recessive lethals per human, predicting 16.0 and 12.2 such mutations, respectively (Fig. 4). Due to this overabundance of strongly deleterious variation in these DFEs, the overall predicted inbreeding load greatly overshoots the observed empirical range. These results become even more extreme for the Kardos et al. (27) model when assuming the original genomic deleterious mutation rate from this paper of U=1.2 (Fig. S5). Under this mutation rate, the Kardos et al. (27) model predicts 31.9 recessive lethals per diploid and a total inbreeding load of 2B=40 (Fig. S5). By contrast, results for the Pérez-Pereira et al. (23) model under the original assumed U=0.4 are somewhat less extreme, though still well outside of the range of empirical estimates (Fig. S5).

In summary, these results demonstrate that the shortcomings of existing deleterious mutation models in terms of their ability to predict empirical estimates of the inbreeding load. These shortcomings motivated our work to develop a new model based on recent papers that directly estimate the DFE for recessive lethal mutations in humans and *Drosophila* (24) and perform DFE inference under non-additive models (26). We believe this model represents the best available deleterious mutation model for humans (and likely other mammals) as it is informed by many sources of information on deleterious mutations, including genetic variation data, empirical inbreeding load estimates, and empirical estimates of segregating recessive lethals (Figs 3 and 4). Notably, our best available model suggests a relatively even partitioning of inbreeding depression in humans due to highly deleterious lethal and semi-lethal mutations and detrimental mutations, which is consistent with studies that suggest a significant fraction of inbreeding depression in mammals being due to deleterious mutations of more modest effect (43, 44). This result implies that, although a substantial portion of the inbreeding load can be purged, it is unlikely to be entirely purged given that negative selection against detrimental mutations may be ineffective in populations of very small size.

**5. The influence of effective population size on inbreeding load**

As discussed above, population size is a critical component influencing the inbreeding load ((17, 23, 27); Fig. S6). This behavior is due to the strong dependence of strongly deleterious and highly recessive ($h < 0.05$) mutations on effective population size (17, 45–47). For example, Kardos et al. (27) showed that the predicted inbreeding load in their model ranged from ~0.5 to ~4 as $N_e$ varied from 25 to 1500 (see Figure S3 in (27)), claiming that this analysis validates their model, as it encompasses the 2B=3.1 estimate from (48). Similarly, Pérez-Pereira et al. (23) claim that their deleterious mutation model is validated by predicting the 2B=12 estimate from (42) when assuming $N_e$=10,000. However, no justification is provided for the assumed effective population sizes assumed in these papers.

To better assess what a reasonable range of equilibrium effective population sizes for mammals, we obtained estimates of equilibrium $N_e$ based on published estimates of genome-wide heterozygosity ($\pi$) from 42 mammal species/populations, including humans. We then estimated equilibrium $N_e$ for each species using the following equation:

$$N_e = \pi/(4*\mu)$$

Given that mutation rate estimates are not available for all species, we assumed $\mu$=1e-8 per site, a mutation rate that is the middle of the range of available estimates (49). Thus, the aim of this analysis is not to precisely estimate equilibrium $N_e$ for each species, but rather to obtain a rough approximation of a median equilibrium $N_e$ in mammals.

From this dataset, we estimate a median equilibrium $N_e$ of 21,875, with a 5% quantile of 3,124 and 95% quantile of 89,703 (Fig. S4). Thus, equilibrium effective population sizes are likely much larger than those assumed in Kardos et al. (27) and Pérez-Pereira et al. (23). For example, the $N_e$ range explored by Kardos et al. (27) of 25 to 1500 is almost entirely within the 1% quantile of this distribution. Moreover, this analysis also demonstrates that humans have an equilibrium $N_e$ that is much more representative for mammals ($N_e$ = 25,936), and that even species with exceptionally low genome-wide diversity, such as the vaquita porpoise, tend to have $N_e$ >> 1000 ($N_e$ = 2,625 for the vaquita).

Several caveats are associated with this analysis. First, as noted above, we consider the same mutation rate of $\mu$=1e-8 for all species, even though some variation is undeniably present. For example, mutation rate estimates in humans are somewhat higher ($\mu$=1.5e-8, implying $N_e$=17,290), whereas the available mutation rate estimate in vaquitas is somewhat lower ($\mu$=5.8e-9, implying $N_e$=4,526). However, a more accurate mutation rate estimate for each species is unlikely to change the conclusion that equilibrium $N_e$ in mammals are generally much larger than those assumed by Kardos et al. (27) and Pérez-Pereira et al. (23). Second, the dataset of genome-wide heterozygosity in mammals is biased towards large, charismatic, and often highly endangered species, which tend to have lower diversity. Specifically, this dataset contains only one rodent (Southeast Asian *Mus musculus*), a species that has by far the highest equilibrium $N_e$=202,266. Moreover, even a broad survey of mammals may be unrepresentative of the range of equilibrium effective population sizes observed in other plant and animal taxa, where $N_e$ estimates are commonly on the order 1e5 to 1e6 (e.g., (15, 38, 39)). Thus, the actual median $N_e$ observed in natural populations is probably far larger than the 21,875 estimated here. Additionally, it should be emphasized that most populations are likely not at equilibrium. For example, although humans are estimated to have an equilibrium $N_e$ on the order of ~20,000, estimates of present-day $N_e$ in humans are often on the order of several hundred thousand (7, 50). Finally, we also note that the model underlying estimation of equilibrium $N_e$ assumes no selection, though the effects of linked selection are known to be widespread across genomes and can greatly contributed to a reduction in diversity relative to what is expected under a neutral model (51). This may contribute to a further underestimation of the true effective population size. Overall, these considerations emphasize that careful modelling of effective population size, ideally based on a demographic model inferred from genomic data, is critical for accurately predicting the inbreeding load in a given species.

**6. Discussion of empirical inbreeding load estimates**

Our analysis makes use of empirical inbreeding load estimates to validate models (Figs 4 and S5). However, not all estimates of the inbreeding load are reliable, which in some cases may limit how meaningful these comparisons are. Obtaining an accurate estimate of the inbreeding load is not trivial: it requires large sample sizes, accurate estimates of the inbreeding coefficient ideally from genomic data, high variance of inbreeding in a population, and a reliable proxy for fitness (41, 52). Studies that combine these elements are rare, leading to wide variance in available estimates. For example, the inbreeding load estimates from 40 captive mammal populations reported in Ralls et al. (37) range from -1.4 to 30.3, with the familiar median of 2B=3.1. Although much of this variance may be explained by small sample sizes and limited variance in inbreeding (37, 52), large confidence intervals for inbreeding load estimates are nevertheless expected even under more ideal conditions. For example, Nietlisbach et al. (41) demonstrated that, even in simulated scenarios with large sample sizes and precise genomics-based estimates of inbreeding, confidence intervals around resulting inbreeding load estimates remain relatively large. Thus, empirical inbreeding load estimates for any given species should be interpreted with care, particularly when sample sizes are small, variance in inbreeding is low, and genomics-based measures of the inbreeding coefficient are not employed.

The meta-analysis presented in Nietlisbach et al. (41) highlighted another crucial issue with existing empirical inbreeding load estimates: many are overestimates due to using biased statistical models. For example, several estimates used in the meta-analysis of O'Grady et al. (42), including the three highest reported estimates of 2B = 13.4, 8.8, and 8.1 were found to be overestimates due to using biased statistical models or unreliable due to issues with the original datasets (41). Moreover, another issue with O'Grady et al. (42) that appears to have been overlooked, but likely also contributes to overestimation of 2B, is their approach of estimating inbreeding load for total fitness by summing separate published estimates from juvenile mortality, survival to sexual maturity, and fecundity. By doing so, this analysis implicitly assumes that these different fitness components are controlled by an entirely non-overlapping set of mutations. However, pleiotropy for mutations underlying traits is known to be widespread (53, 54), suggesting that this independence assumption does not hold. In other words, the approach employed by O'Grady et al. (42) likely results in double or triple counting the effects of deleterious mutations contributing to inbreeding load, further contributing to upward bias in their estimate of 2B=12. For these reasons as well as those outlined by Nietlisbach et al. (41), the 2B=12 estimate for wild populations from O'Grady et al. (42) is likely to greatly overestimate the true inbreeding load and should therefore not be employed by simulation studies. The median estimate from Nietlisbach et al. (41) of 2B=4.5 is much more likely to be reflective of the true inbreeding load for survival to sexual maturity in wild vertebrates, though it should be noted that this estimate is based on a relatively few species, nearly all of which are birds.

In summary, although empirical inbreeding load estimates should be interpreted with care, we suggest that careful meta-analyses of available inbreeding load estimates, such as that of Nietlisbach et al. (41), can still provide a useful point of comparison. The relatively good agreement with the median estimate of 2B=4.5 from Nietlisbach et al. (41) with our model prediction of 2B=6.2 for humans helps validate our model parameters, which are derived from very different sources of information. Importantly, we do not necessarily expect that our inbreeding load predictions will perfectly agree with median estimates from Ralls et al. (37) and Nietlisbach et al. (41), as it is expected that the actual inbreeding load for a given species will vary. Rather, these estimates serve to place a rough bounds on what might be expected for the total inbreeding load in humans, given the limitations with the human estimate from Bittles & Neel (40) discussed above. Moreover, it is encouraging that the predicted inbreeding load from our best available model due to lethals and semi-lethals of 2B=3.1 is in good agreement with the estimates based on juvenile mortality Bittles & Neel ((40); 2B=1.4) and Ralls et al. ((37);

2B=3.1), which might be expected if it is assumed that juvenile mortality is largely a consequence of lethal and semi-lethal mutations. Finally, our results also suggest that the inbreeding load due to detrimentals may be greatly underestimated in humans, which may be expected given the challenges of quantifying fitness in humans.

### 7. Supplementary Figures and Tables

**Table S1: Methods for detecting and quantifying different types of deleterious mutations.** For each mutation type, the table lists what methods are best suited for quantifying such mutations, the mutation rate per diploid genome of such mutations (U), the mean selective coefficient of such mutation (s), and the approximate % of deleterious mutations that fall in each category. Note that lethal and semi-lethal mutations are likely a subset of nonsynonymous mutations, though are better detected by experimental methods than genetic-variation-based methods. Thus, these values imply an overall U = ~2.5 for deleterious mutations. Note that all values are approximate and intended to convey order of magnitude, and are likely to vary across species.

| Deleterious mutation type | Detection method | U | Mean *s* | % of all deleterious mutations | References |
|---|---|---|---|---|---|
| Lethal and semi-lethal mutations | experimental | 0.01-0.05 | -0.2 | 1 | (3, 22, 55) |
| Nonsynonymous mutations | genetic variation data | 0.2-0.6 | -0.01 | 20 | (7–9, 28, 29, 31, 32) |
| Mutations in conserved non-coding regions | evolutionary constraint; genetic variation data | 1-2 | -0.001 | 80 | (11, 28, 29, 56, 57) |

**Table S2: Comparison of predicted nonsynonymous SFS from four DFE and dominance models to SFS from 432 unrelated European individuals in 1000G.**

| SFS | % singletons | % common variants (>5% frequency) |
|---|---|---|
| Empirical 1000G | 56.8 | 13.3 |
| Kyriazis et al. 2021 | 53.9 | 10.6 |
| Kardos et al. 2021 | 76.2 | 2.9 |
| Pérez-Pereira et al. 2022 | 72.8 | 5.4 |
| This paper | 57.3 | 10.4 |

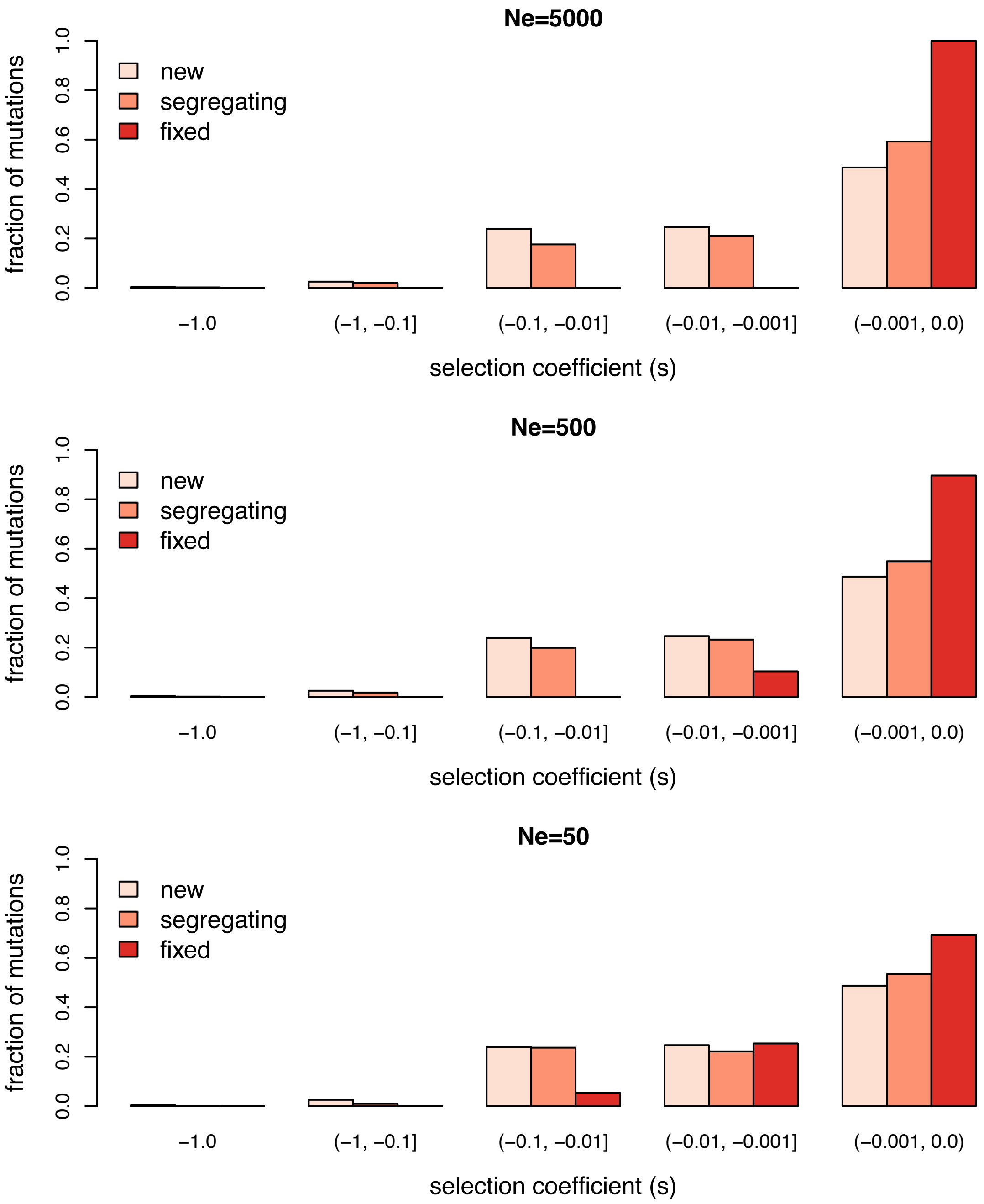


**Figure S1: Comparison of the distribution of *s* for new, segregating, and fixed mutations when $N_e$=50, $N_e$=500, and $N_e$=5000 at equilibrium**. For this analysis, we simulated using our best available model along with human genomic parameters, as described in Supplemental Appendix 3. Note the differences between the distribution of *s* for new mutations and distribution for segregating and fixed mutations in each case.

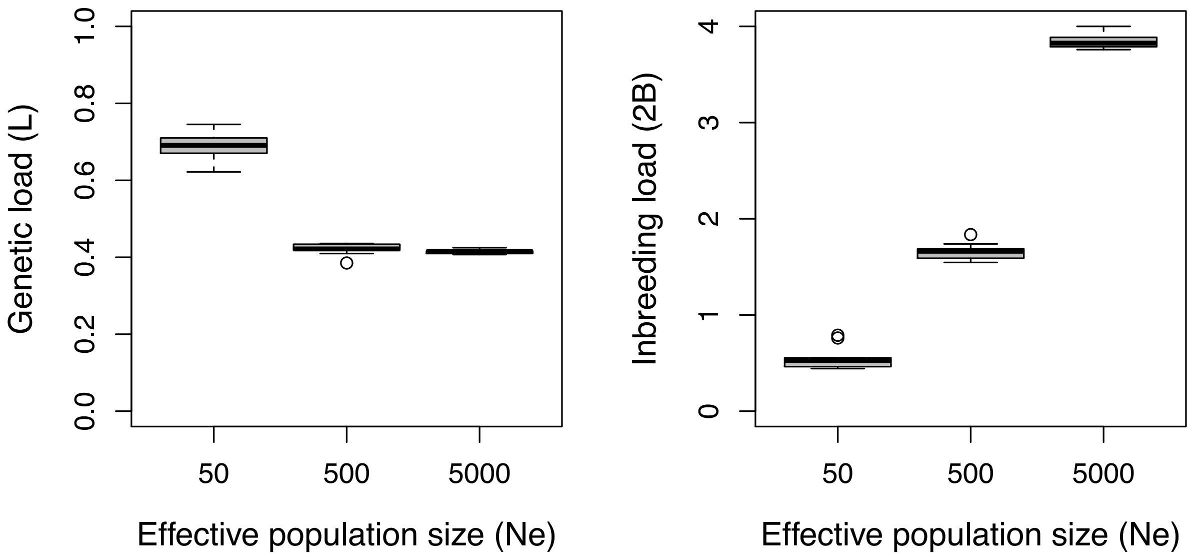


**Figure S2: Boxplots of genetic load and inbreeding load from a simulated example.** For this analysis, we used our 'best available' deleterious mutation parameters along with human genomic parameters, as described in Supplemental Appendix 3. Note that both genetic load and inbreeding load depend on $N_e$, but this relationship is not always linear.

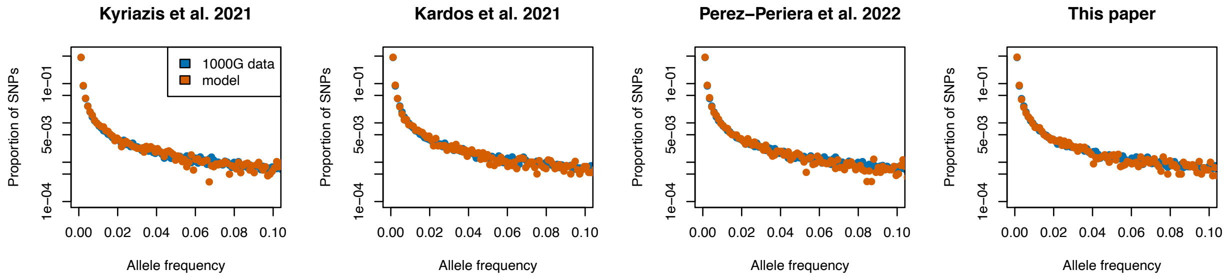


**Figure S3: Predicted proportional synonymous SFS for simulations shown in Figure 4 in comparison to synonymous SFS from 864 haploids in 1000G dataset.** Simulations were conducted using a demographic model inferred by Kim et al. 2017 using the 1000G synonymous SFS, thus the fit of the simulated synonymous SFS to the data is expected in all cases. Simulation results are based on ~8.1 Mb of simulated coding sequence.

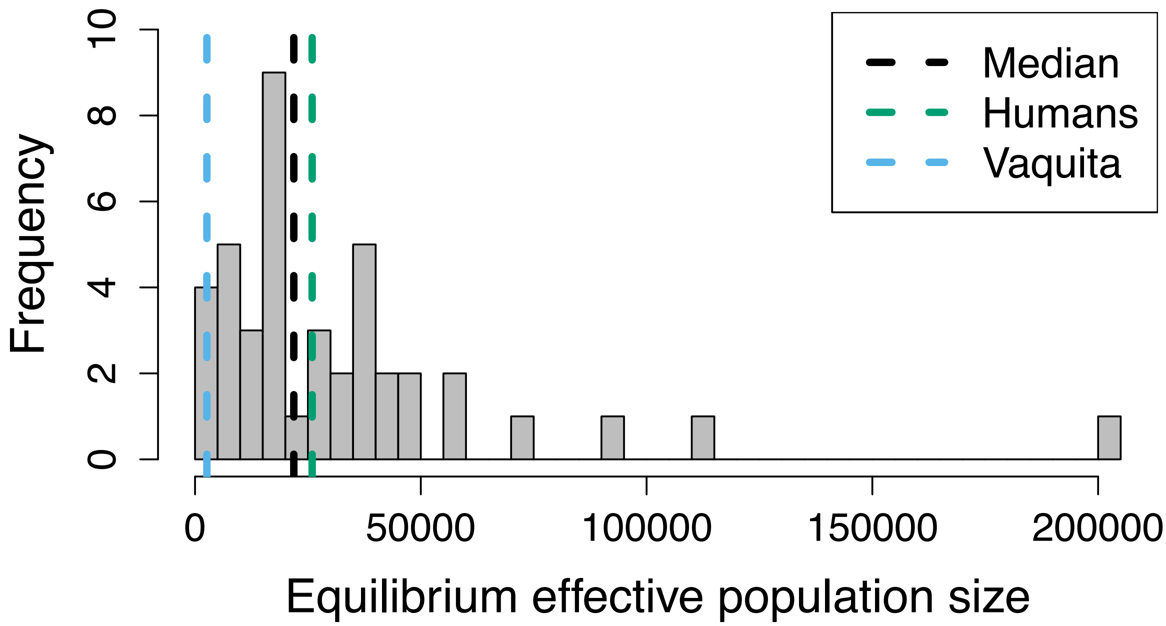


**Figure S4: Histogram of equilibrium effective population size based on genome-wide diversity estimates in a sample of 42 mammal species/populations.** Dashed vertical lines indicate median estimate (Ne = 21,875), estimate for humans (Ne = 25,936), and estimate for the vaquita (Ne = 2,625). For all species, we assume a mutation rate of 1e-8 per site.

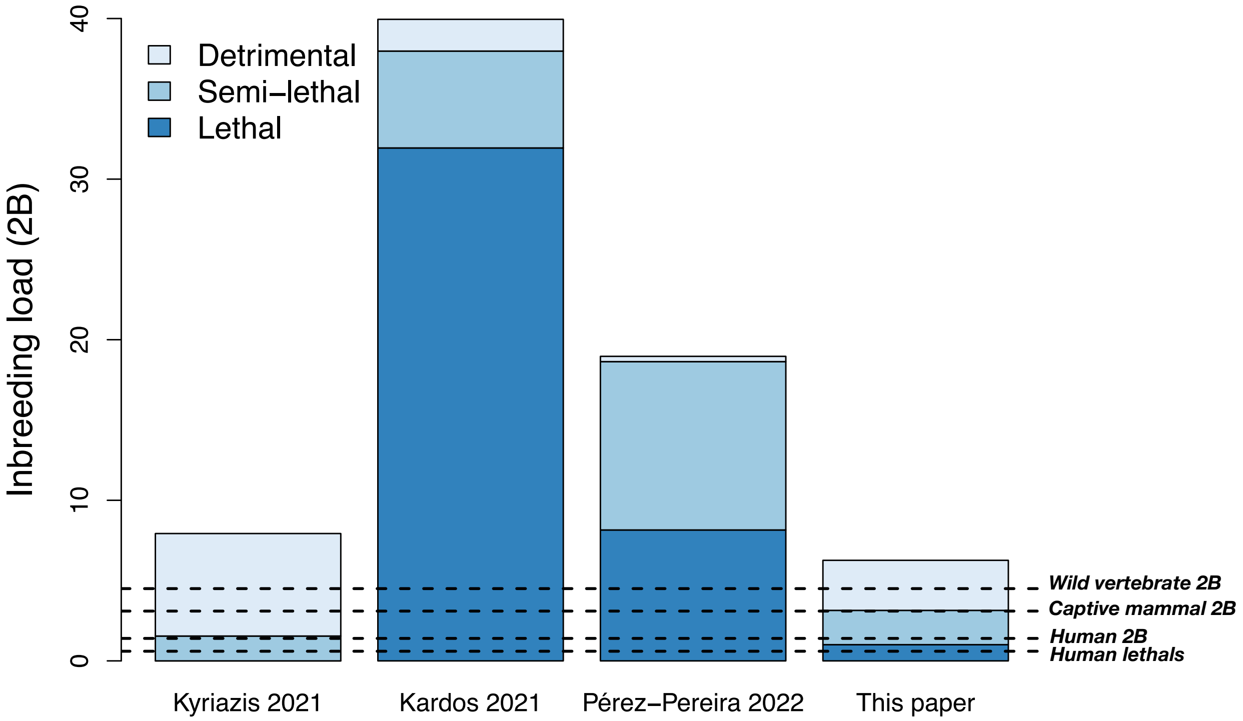


**Figure S5: Inbreeding load partitioning under various DFE and dominance models using demographic and genomic parameters for humans, plotted with the original genomic deleterious mutation rates assumed by each study (U=0.42 for Kyriazis et al.** (17)**, U=1.2 for Kardos et al.** (27)**, U=0.4 for Pérez-Pereira et al.** (23)**, and U=0.63 for 'best available' model from this paper).** Colors depict contribution of inbreeding load from each class of deleterious mutations, with the total height of each bar representing the total predicted inbreeding load (2B). Detrimentals are here defined as mutations with $s$ > -0.1, semi-lethals as mutations with -0.99 < $s$ <= -0.1, and lethals as mutations with $s$ < -0.99. Dashed lines show estimated of number of lethals per diploid human from Gao et al. (25)("human lethals"), inbreeding load estimate for humans from Bittles & Neel (40)("human 2B"), and estimate of average inbreeding load for vertebrates from Nietlisbach et al. (41) ("vertebrate 2B"). Note that predicted inbreeding load partitioning under the model proposed in this paper agrees well with empirical estimates, whereas predicted inbreeding load partitioning from other models do not.

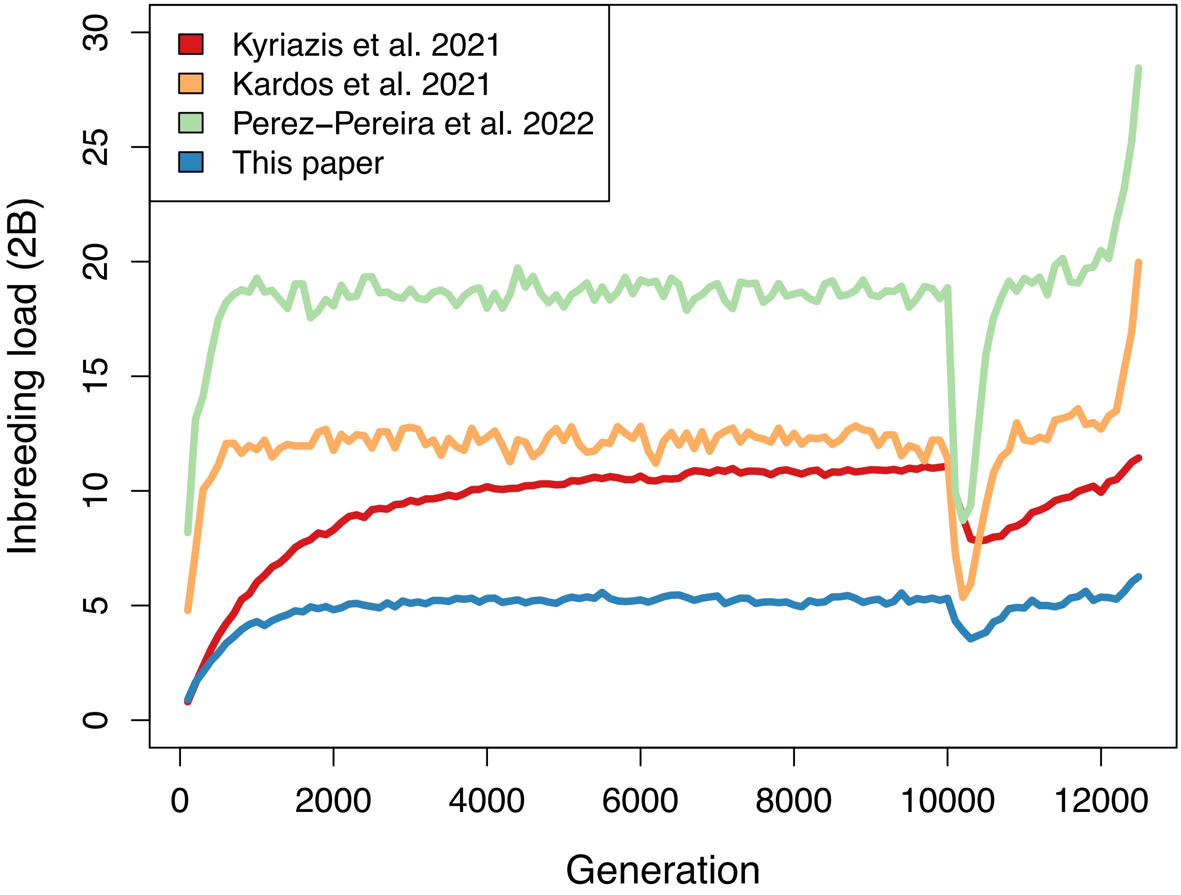


**Figure S6: Inbreeding load dynamics over time during simulations. Plotted is inbreeding load (2B) for each DFE and dominance model under the demographic model inferred by Kim et al.** (7) **with a 10,000 generation burn-in duration.** Note that all models reach equilibrium by ~8,000 generations, though increase to equilibrium and increase during exponential growth is much faster for the Kardos et al. (27) and Pérez-Pereira et al. (23) models due to the high fraction of recessive lethals in these models. These dynamics are also reflected during the out-of-Africa bottleneck, where purging is far greater for the Kardos et al. (27) and Pérez-Pereira et al. (23) models.
